# Reproductive isolation emerges from coordinated barriers during mating system divergence in plants

**DOI:** 10.64898/2026.04.26.720968

**Authors:** Ana García-Muñoz, Camilo Ferrón, Enrica Olivieri, Mohamed Abdelaziz, A. Jesús Muñoz-Pajares

**Affiliations:** BioChange network, Departamento de Genética, Universidad de Granada, Granada, Spain; Departamento de Biología Vegetal y Ecología, Universidad de Sevilla, Sevilla, Spain; Área de Biodiversidad y Conservación, Departamento de Biología y Geología, Física y Química Inorgánica, Universidad Rey Juan Carlos, Móstoles, Spain

**Keywords:** Ecological speciation, Mating system evolution, Pollinator-mediated isolation, Pre- and post-pollination barriers, Hybridization, *Erysimum* L., Magic traits

## Abstract

**Background and Aims:** Understanding how reproductive barriers combine to restrict gene flow remains a central challenge in speciation research. Although reproductive isolation is inherently a composite process, most empirical studies have focused on individual barriers in isolation, limiting our ability to capture their joint effects particularly in systems undergoing evolutionary transitions such as shifts in mating system.

**Methods:** Here, we provide a comprehensive, life-cycle-wide quantification of reproductive isolation between two closely related species of the *Erysimum incanum* complex that differ strikingly in mating system: the predominantly selfing *E. incanum* and the outcrossing *E. wilczekianum*.

**Key Results:** By integrating ecological, phenological, behavioural, and post-pollination components, we show that total reproductive isolation is nearly complete (T≈0.999), but overwhelmingly driven by pre-pollination barriers. Ecogeographical differentiation and, most prominently, pollinator-mediated isolation dominate, with pollinators exhibiting a strong bias toward *E. wilczekianum*. Floral traits linked to mating system divergence, particularly flower size, emerge as key drivers of assortative mating, supporting their role as “magic traits” coupling ecological divergence with reproductive isolation. In contrast, post-pollination barriers are weaker but strongly asymmetric. Hybrid seed formation is largely prevented when *E. wilczekianum* acts as the maternal parent, consistent with expectations from mating system differences, whereas reciprocal crosses are relatively successful. Despite reduced germination, hybrids display enhanced growth and no evidence of hybrid breakdown, suggesting that intrinsic incompatibilities remain incomplete.

**Conclusions:** These findings reveal that mating system divergence restructures the entire architecture of reproductive isolation rather than acting as a single barrier. More broadly, our results highlight that early-stage speciation can be driven by coordinated shifts in ecological and reproductive traits, emphasizing the need for integrative approaches to fully understand how barriers interact to generate species boundaries.

## INTRODUCTION

The speciation process relies on the accumulation of genetic differences among populations, ultimately driven by the emergence of reproductive barriers that reduce or prevent gene flow between them (Mayr, 1947). However, reproductive isolation is not the result of a single mechanism, but rather the outcome of multiple barriers acting across the life cycle, whose combined effects determine the extent of gene flow reduction (Ramsey et al., 2003; Sobel and Chen 2014). Despite this, most studies have focused on individual barriers in isolation, overlooking how their interaction shapes the speciation process. This limitation becomes particularly relevant in systems undergoing evolutionary transitions, such as shifts in mating system, where multiple barriers may change simultaneously and in a coordinated manner. Understanding how these barriers integrate and contribute to total reproductive isolation is therefore essential to uncover the mechanisms underlying speciation. When these barriers arise as a consequence of adaptation to different environments, speciation can proceed through ecological divergence, a process widely recognized as ecological speciation (Coyne and Orr, 1998; Nosil, 2012).

Within this framework, identifying traits that simultaneously influence ecological divergence, and reproductive isolation becomes particularly relevant. In many cases, speciation involves multiple traits, some directly contributing to reproductive isolation and others primarily shaped by ecological selection. However, certain traits can simultaneously mediate both processes. These so-called *magic traits* link ecological divergence and reproductive isolation, thereby facilitating speciation even in the presence of gene flow (Servedio and Kopp, 2012). Although most commonly described in animals, floral traits such as size, colour or morphology may act as magic traits in plants, as they influence both adaptation to pollinator environments and patterns of assortative mating (Servedio et al., 2011; Haller et al., 2012).

Floral attractiveness has a strong influence on the frequency and diversity of pollinator visits (Stanton et al., 1986; Erickson and Pessoa, 2022), which constitute the primary vectors of gene flow among plant individuals (Handel, 1983; Deynze et al., 2005; Hoyle and Cresswell, 2007; Wang et al., 2021). Consequently, divergence in floral traits can alter pollinator assemblages and behavior, promoting assortative mating and contributing to reproductive isolation. Despite this, gene flow often persists between incipient species. Indeed, hybridization is common in plants, with a substantial proportion of species forming hybrids in nature (Mallet, 2007). This prevalence highlights the difficulty of applying strict species boundaries based solely on reproductive isolation and underscores the importance of quantifying gene flow during the speciation process.

Reproductive isolation is not a single mechanism but rather the cumulative outcome of multiple barriers acting throughout the life cycle (Ramsey et al., 2003; Nosil, 2007; Lowry et al., 2008a; Matsubayashi and Katakura, 2009; Jiménez-López et al., 2023). These barriers are typically classified according to whether they act before or after pollination, commonly referred to as pre- and post-pollination barriers (Ramsey et al., 2003; Lowry et al., 2008a; Baack et al., 2015; Jiménez-López et al., 2023). The relative strength and interaction of these barriers determine both the magnitude of gene flow reduction and the stage of divergence between taxa. In general, pre-pollination barriers tend to dominate in early stages of speciation, whereas post-pollination barriers become more prominent as divergence increases (Hendry et al., 2007; Nosil, 2012).

Pre-pollination barriers include ecological and behavioral processes that limit the probability of mating. Ecogeographical isolation arises from differences in habitat preference and spatial distribution, reducing the likelihood of encounter between potential mates (Honnay et al., 2002; Sobel et al., 2010). These patterns can be evaluated through species distribution modelling, which provides a robust framework to estimate habitat overlap and potential contact zones (Peterson, 2003; Kozak et al., 2008). Phenological isolation, in turn, occurs when temporal mismatches in flowering reduce mating opportunities, even when species co-occur (Primack, 1985; Husband and Schemske, 2000). Finally, pollinator-mediated isolation depends on the degree of pollinator fidelity and the extent to which floral traits influence pollinator behaviour (Grant, 1949).

Post-pollination barriers act after pollen deposition and include processes affecting fertilization success and hybrid performance. The interaction between pollen and stigma represents a critical checkpoint, often influenced by the mating system of the parental species and associated molecular mechanisms (Lowry et al., 2008a; Sambatti et al., 2012). Differences in mating system, particularly between selfing and outcrossing lineages, can generate strong asymmetries in reproductive isolation, partly due to differences in parental conflict and pollen–pistil interactions. Beyond fertilization, hybrid viability and fertility determine the ultimate success of gene flow, with outcomes ranging from hybrid vigour to hybrid breakdown across generations (East, 1936; Templeton et al., 1986).

Despite the importance of integrating all components of reproductive isolation, most empirical studies have focused primarily on pre-pollination barriers (Ferriol et al., 2009; Nosil, 2012; Matsumoto et al., 2019; Cardona et al., 2020 but see Jiménez-López et al., 2023). As a consequence, post-pollination processes are often underestimated, particularly in systems at early stages of divergence. Moreover, many studies are restricted to a limited number of taxa or do not quantify the relative contribution of individual barriers to total reproductive isolation (Baack et al., 2015). However, reproductive barriers do not act independently; their interaction can either reinforce or constrain each other, influencing the trajectory of speciation (Nosil and Yukilevich, 2008; Agrawal et al., 2011).

Here, we investigate the extent of reproductive isolation between two closely related species within the *Erysimum incanum* species complex that exhibit contrasting mating systems. We quantify total reproductive isolation and the relative contribution of multiple barriers acting across the life cycle. Specifically, we evaluate pre-pollination barriers including ecogeographical separation, flowering phenology and pollinator preferences, as well as post-pollination barriers from pollen–stigma interactions to hybrid viability and fertility across generations. By integrating these components, we aim to provide a comprehensive assessment of the mechanisms underlying reproductive isolation and to better understand the role of mating system divergence in plant speciation.

## MATERIAL AND METHODS

### Study system

We focused on two species of the genus *Erysimum* L., one of the most diverse genera within Brassicaceae, distributed across Eurasia, North Africa, and the Americas (Al-Shehbaz et al., 2006). These species differ markedly in both mating system and floral traits. *Erysimum incanum* is predominantly selfing and exhibits the typical features of the selfing syndrome (Sicard and Lenhard, 2011), including small flowers, reduced nectar and pollen production, and a low pollen–ovule (P/O) ratio. Three ploidy levels have been described in this species—diploid, tetraploid, and hexaploid (Nieto Feliner, 2014; Chapter 1). Diploid populations occur in the Rif and Pyrenees Mountains, tetraploids are found in southwestern Iberia and the Middle Atlas, and hexaploids are restricted to the High and Middle Atlas Mountains (Fennane and Ibn-Tattou, 1999). In contrast, *Erysimum wilczekianum* is an outcrossing species pollinated by insects, characterized by larger flowers and strong inbreeding depression (García-Muñoz, 2023). Its distribution is restricted to a narrow region in Morocco, where it occurs near tetraploid populations of E. incanum (Fennane and Ibn-Tattou, 1999). Phylogenetic analyses indicate that *E. wilczekianum* belongs to an outcrossing clade nested within a predominantly selfing lineage (Abdelaziz et al., in prep.; García-Muñoz, 2023), challenging the traditional view of selfing as an evolutionary dead end. This contrast in mating systems and floral traits makes this species pair an excellent system to investigate ecological divergence and reproductive isolation driven by traits with dual ecological and reproductive roles.

### Reproductive isolation estimation

We quantified reproductive isolation across the full life cycle of both taxa using a combination of field observations and controlled greenhouse experiments. Total reproductive isolation and the relative contribution of each barrier were estimated following Ramsey et al. (2003) and Sobel and Chen (2014).

### Pre-pollination reproductive barriers

#### Ecogeographical isolation

We found a minimal distance of 500 m between *E. wilczekianum* and the closest *E. incanum* population, which are both tetraploids (Fennane and Ibn-Tattou 1999). This distance could be based on differences in habitat preferences exhibited by each species. For this reason and to cope with possible mapping distribution dismissal, we decided to test the differences in habitat preferences and the potential overlapping degree by SDM. We mapped the spatial distribution of four tetraploid *E. incanum* populations and two *E. wilczekianum* populations inhabiting Morocco.

Based on our own records, we constructed the distribution model of both species in Maxent v.3.4.4 (Phillip, 2010) using the principle of maximum entropy (Guisan and Thuiller, 2005). We used bioclimatic variables from Worldclim v.2 dataset (www.wordclim.org; Fick and Hijmans, 2017) at 1 km^2^ resolution. After discarding variables with high collinearity (Guisan et al., 2017), we focused on the average precipitation (VIF = 1.84), average temperature (VIF = 3.03) and two other bioclimatic variables related to precipitation seasonality and the mean diurnal range of temperature (VIF = 3.26 and VIF = 1.87, respectively) which have been demonstrated to be key for determining plant distribution by other works (Dixon and Busch, 2017; Duffy and Jacquemyn, 2019). We also considered elevation and geological variables from ISRIC-WISE v3.1 database (Batjes, 2012). Distribution model was built on 50 Maxent replicates selecting the cloglog format output. We maintained standard default settings (converge threshold = 0.00001, iterations = 500, default prevalence = 0.5) and used the equal training specificity and sensitivity (ETSS) approach in order to classify accurately suitable or unsuitable sites. This threshold represents the umbral at which predictions about suitability or unsuitability habitats have an equal probability of being correct (Freeman and Moisen, 2008).

To assess the differences in niche preferences we compared the species distribution models using the Schoener’s D index (Schoener, 1970), which is an indicator of habitat overlapping between the species. Specifically, we used the niche identity test implemented in the R package ENMTools v.1.1.5 (Warren et al., 2025) in R v4.2.3 (R Core Team, 2023). Niche identity test uses the Schoener’s D index to evaluate the probability of two species niches to be identical. This metric ranges from 0 when niches are not overlapping, to 1 when niches of both species are completely overlapped (Schoener and Gorman, 1968; Püts et al., 2020). This index has been used in studies of reproductive isolation in plants as the ecogeographical barrier (Glennon et al., 2012; Willis and Donohue, 2017) as:

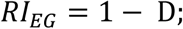

where D is Schoener′s D index. We compared the empirical value of the D index from our data with the null distribution generated by 100 randomly permuted species distribution models in order to test the existence of significant differences in habitat preferences between the two species.

### Phenological isolation

According to our personal observations, the flowering for both species overlaps in the wild, with *E. wilczekianum* starting to flower earlier. To remove environmental effects, we tested the phenological barrier in common greenhouse conditions. We sowed seeds from two populations of *E. wilczekianum* and from the closest population of *E. incanum*. We checked germinated plants daily and, once the first flower opened in one of the individuals, we monitored plants every two days during a month. Specifically, we recorded the number of plants showing at least one open flower and the number of open flowers per plant. We also noted the date of flowering termination for each species and estimated the phenological barrier by the degree of overlapped flowering following the method proposal by (Husband and Sabara, 2004):

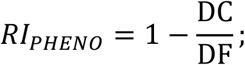

where DC is the number of days co-flowering in both species and DF is the number of total number of days flowering in a single species.

### Pollinator preference isolation

We tested the reproductive barrier driven by pollinator preferences by setting an experimental array with 30 potted plants (15 per species) separated from each other by 0.5 m and randomly arranged in a rectangle (5×6 pots). Before starting the observations, we measured floral traits (petal length, corolla diameter and corolla tube length) from one flower of each individual of the array and counted the number of flowers in anthesis at the moment of the observations in order to test the effect of specific trait values on visit frequency. The experiment was performed in two sites, corresponding to the habitat of *E. incanum* and the habitat of *E. wilczekianum.* We repeated the observations twice in each site, moving the pots randomly every day. Two observers noted flights that insects performed between plants from the same or different species during observation periods of 50 minutes between 10.30 am and 17.30 pm, local time. We only noted the flights where pollinators touched the reproductive part of the flower, considering the pollen transfer as effective. We obtained a symmetric measurement of isolation between both species:

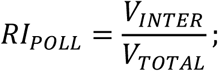

where V_INTER_ is the number of interspecific visits and V_TOTAL_ is the number of total visits. Finally, we compared the amount of visits between the two sampe sites by ANOVA analyses in order to detect whether the frequency of visits is influenced by the local pollinator assemblage. We assessed if pollinators showed preferences by the traits exhibited in each species by Pearson’s product-moment correlations.

### Post-pollination reproductive barriers

#### Hybrid seeds production

We estimated the first post-pollination barrier to gene flow comparing the seed production from intraspecific and interspecific crosses. These crosses were performed in both directions, that is, *E. incanum* and *E. wilczekianum* acting both as pollen donor and receptor. To perform hybrid crosses in *E. incanum*, we carefully emasculated flowers before opening in the plant receptor to avoid the anther-rubbing mechanisms described in this species (Abdelaziz et al., 2019). Then, we added pollen from *E. wilczekianum* plants. Since *E. incanum* is predominantly selfing according to previous studies (García-Muñoz et al., 2025) auto-pollinated crosses served as control in *E. incanum.* Following the same rationale, to perform hybrid crosses in *E. wilczekianum* we emasculated flowers before opening and added pollen from *E. incanum* plants. Because *E. wilczekianum* exhibits high values of inbreeding depression (García-Muñoz et al., 2023), we performed cross-pollinated crosses between different individuals to obtain the seed production from intraspecific crosses which served as control. In all cases, each crossed flower was labeled to identify the type of treatment once the plant life cycle was finished. When it occurred, fruits were recollected to count the number of viable seeds, aborted seeds and non fertilized ovules. The sum of these three parameters resulted in the amount of produced ovules per flower. The seedset was the number of viable seeds divided by the total amount of ovules. In total, we conducted 162 hybrid crosses in *E. incanum* and 131 hybrid crosses in *E. wilczekianum*. We used 1204 and 954 control crosses in *E. incanum* and *E. wilczekianum*, respectively. All these crosses involved 121 *E. incanum* and 255 *E. wilczekianum* plants. We estimated reproductive barrier due to hybrid seed formation applying the method proposed by (Lowry et al., 2008b) to each species:

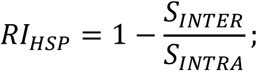

where S_INTER_ is the seedset from interspecific crosses and S_INTRA_ is the seedset from intraspecific crosses.

### Viability and fertility of the hybrids

Finally, we sowed a subset of 300 hybrid seeds resulting from *E. incanum* and 300 seeds from each parental species. We only selected viable seeds which are easily differentiated from aborted seeds by the color and consistency. We estimated the hybrid viability by comparing its germination with the germination of the parental seeds. Since we did not sow seeds of hybrids from *E. wilczekianum*, we compared the hybrid germination of *E. incanum* hybrids with both parental species. Germination was measured as the number of germinated seeds that reach the rosette size by the total of seeds that were sowed in each pot. Here, the germination was a proxy of survival because mortality in these plants is not common once the plants become rosettes. The reproductive barrier due to the viability of first hybrid generation (F1) was estimated as:

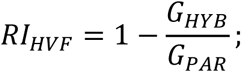

where G_HYB_ is the proportion of germinated hybrid plants and G_PAR_ is the proportion of germinated plants from each parental.

Additionally, we estimated the growth ratio counting the leaves. When plants started to bloom, we measured floral size traits (petal length, corolla diameter and corolla tube length, described above). In addition, we measured nectar production in order to obtain a more accurate picture of differences in floral phenotype between hybrid individuals and parental species. Nectar was measured in a flower per plant measuring nectar amount using glass Drummond® capillary 0.5 µL.

Hybrid fertility was estimated as the seedset produced by these plants. For that, we counted the number of seeds, aborts and unfertilized ovules in four fruits per plant. We did not manipulate flowers, allowing for autonomous self-pollination because the phenotype was similar to *E. incanum* regarding flower size so we considered that the mating system would be similar. The barrier due to hybrid fertility was calculated only for the hybrids from *E. incanum* due to the unbalanced production of hybrids from each species as pollen receptors. The barrier due to the fertility exhibited by the first hybrid generation was calculated as:

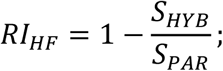

where S_HYB_ is the seedset from hybrid plants and S_PAR_ is the seedset from *E. incanum* parental plants.

We obtained a second generation of hybrids sowing a total of 560 seeds produced by self-pollinated fruits from the first generation of hybrids. We followed the methodology to compare the viability in this second hybrid generation (F2) comparing again with *E. incanum* individuals by:

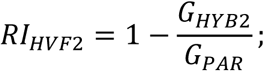

where G_HYB2_ is the proportion of germinated plants from the second hybrid generation and G_PAR_ is the proportion of germinated plants from *E. incanum* parental plants.

### Absolute and relative contribution of reproductive barriers

Following the methods proposed by Coyne and Orr (1998) and Ramsey et al. (2003), we also calculated the absolute and relative contribution of each barrier and the total reproductive isolation. The absolute contribution (AC) of a barrier to the reproductive isolation is the gene flow removed that was not eliminated by previous barriers and can be estimated as:

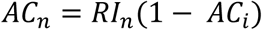

In this notation *n* takes values from 1 to 5 to represent barriers due to ecogeography, phenology, pollinators, hybrid seeds production and hybrids viability and fertility, respectively and *i* refers to the previous barriers. Thus, the absolute contribution of the ecogeography is equal to the estimated reproductive isolation (RI ecogeograhical) whereas for pollinators the absolute contributions of ecogeography and phenology must be subtracted. When the absolute contribution of every barrier has been estimated, it is possible to quantify the total reproductive isolation (T) by summing these values as follows:

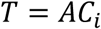

This total reproductive isolation T varies from 0 (complete lack of isolation) to 1 (complete isolation between species). Finally, we can estimate the relative contribution of each barrier (RC_n_) to the total reproductive isolation (T) as:

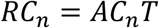

with *n* taking values from 1 to 5 to represent the different barriers as detailed for AC_n_. All these parameters were estimated for both *E. incanum* and *E. wilczekianum* in order to consider the asymmetries in the barriers to gene flow.

## RESULTS

### Total reproductive isolation

The strength of the total reproductive isolation and the absolute and relative component contributions of the different barriers are summarized in Table 1. The species *E. incanum* and *E. wilczekianum* experimented a similar and high total reproductive isolation, with values of T = 0.9988 and 0.9994, respectively. These values were mainly due to pre-pollination barriers such as ecogeographical and pollinator isolation. Even though pre-pollination barriers are high and symmetric, we found a strong asymmetry in post-pollination barriers involving the hybrid seed formation.

**Table 1.** Summary of the individual reproductive barriers, shown by each species and the absolute (AC) and relative (RC) contribution of each barrier to total reproductive isolation (T).

| Reproductive barrier | Components of reproductive isolation (RI) |  | Absolute contribution to total isolation (AC) |  | Relative contribution to total isolation (RC) (%) |  |
| --- | --- | --- | --- | --- | --- | --- |
|  | *E. incanum* | *E. wilczekianum* | *E. incanum* | *E. wilczekianum* | *E. incanum* | *E. wilczekianum* |
| Ecogeographical isolation | 0.3979 | 0.3979 | 0.3979 | 0.3979 | 39.8458 | 39.8227 |
| Floral phenology | 0.2711 | 0.0752 | 0.1633 | 0.04527 | 16.3443 | 4.5306 |
| Pollinator preference | 0.9858 | 0.9858 | 0.4325 | 0.5488 | 43.2985 | 54.9093 |
| Hybrid seed formation | 0.1430 | 0.9260 | 0.0009 | 0.0073 | 0.0892 | 0.7314 |
| H<sub>1</sub> viability | 0.4781 | 0.0410 | 0.0026 | <0.0001 | 0.2563 | 0.0024 |
| Hybrid fertility | 0.1735 | - | 0.0005 | - | 0.0480 | - |
| H<sub>2</sub> viability | 0.5010 | - | 0.0012 | - | 0.1162 | - |
| **Total isolation (T)** |  |  | **0.9988** | **0.9994** |  |  |

### Pre-pollination reproductive barriers Ecogeographical isolation

We used the Maxent approach based on the ETSS threshold that resulted in constructing reliable distribution models for both species according to the obtained parameters. One of these indicators of model quality is the area under receiver operator curve (AUC) which assesses the model fit by plotting the proportion of predicted presences and the proportion of predicted absences (Peterson, 2003; Phillips and Others, 2005). It ranges from 1 (the model is optimum predicting the habitat suitability) to 0.5 (the model is inaccurate in estimating the habitat suitability). The AUC that we obtained from our species distribution models were 0.820 for *E. incanum* and AUC = 0.857 for *E. wilczekianum* species. Error estimates are produced by the 95% confidence intervals on the ETSS threshold values. Distribution range is quite different among species because *E. incanum* shows a broader niche compared to *E. wilczekianum* (Figure 2). The identity test indicated substantial niche overlap (Schoener’s D = 0.6021), although the niches were significantly different from each other (p-value = 0.029). Based on the empirical D index, the reproductive barrier due to ecogeographical differences was 0.3979.

**Figure 1.**
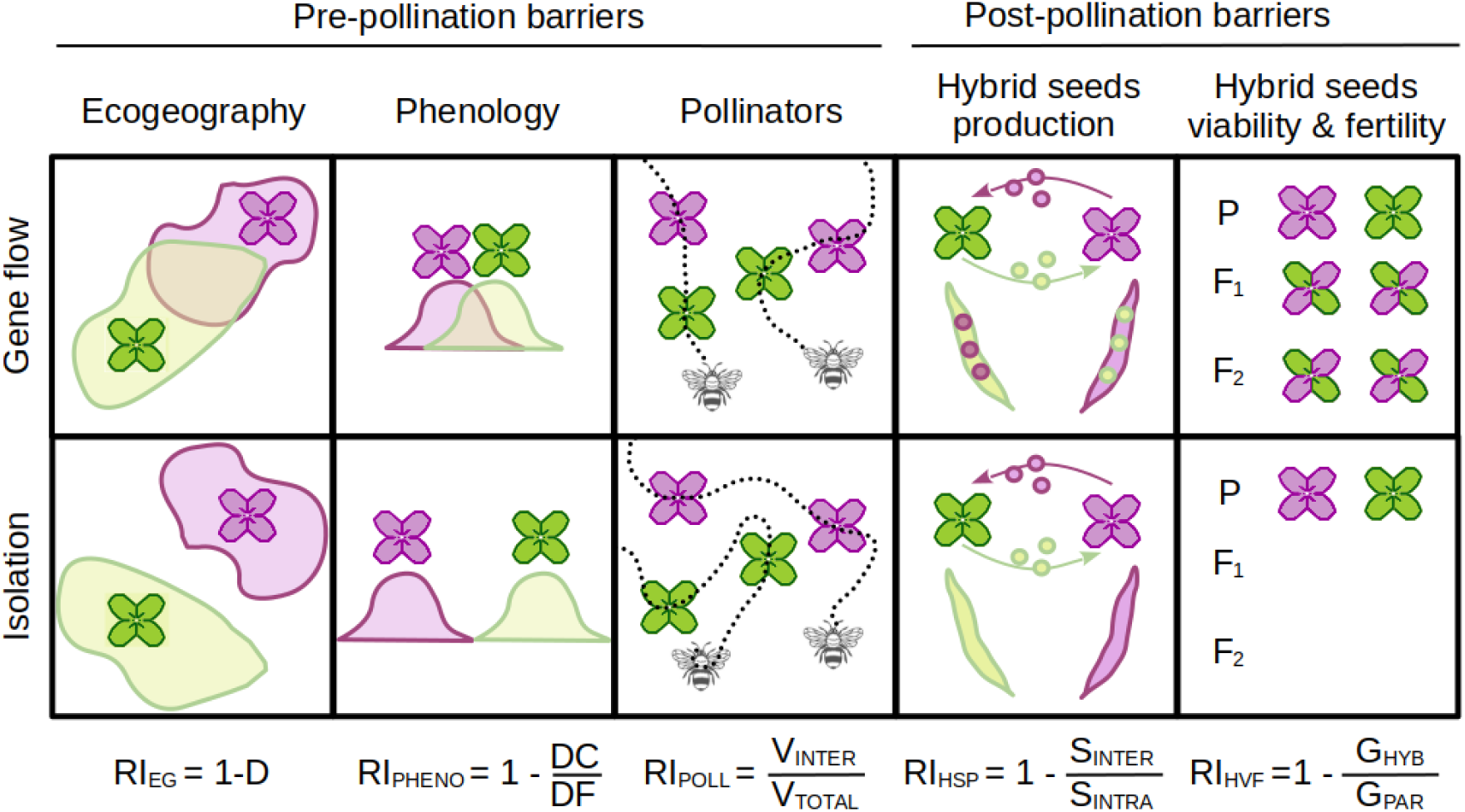
Summary of the expected outcomes for every reproductive barrier under the scenarios of gene flow and reproductive isolation. Equations used to quantify individual reproductive isolation barriers (RI) are also shown (D: Schoener’s D index; DC: days co-flowering; DF: days flowering; V_INTER_: Interspecific visits; V_TOTAL_: total visits; S_INTER_: Interspecific seedset; S_INTRA_: Intraspecific seedset; G_HYB_: Germinated hybrids; G_PAR_: Germinated parentals).

**Figure 2.**
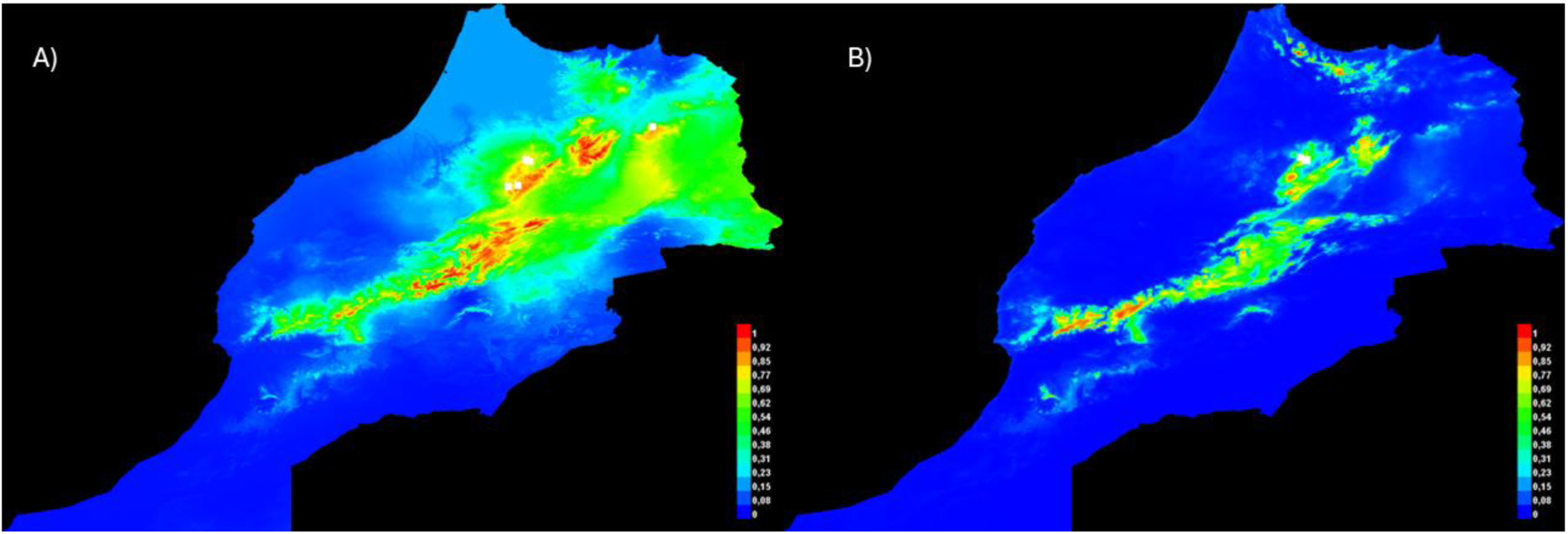
Potential distribution of (A) *E. incanum* and (B) *E. wilczekianum* calculated by Maxent. Colors represent the likelihood of the species occupation. Red zones predict the presence of the species with complete likelihood and blue zones indicate the no existence of the species. White points indicate the real presence of the species.

### Phenological isolation

Plants of *E. wilczekianum* started flowering in early February (7^th^ February was the date of the first flower opening) and first flowers of *E. incanum* opened just a week later (14^th^ February). The two species shared a part of the flowering period but *E. wilczekianum* finished blooming earlier than *E. incanum* (12^th^ May), which kept producing new flowers until 8^th^ June. Although the flowering period was shorter, *E. wilczekianum* showed a greater amount of simultaneously open flowers per plant (Fig. 3). In total, *E. incanum* maintained its flowers open for 118 days while *E. wilczekianum* showed open flowers for 93 days. Flowering overlap between the two species occurred during a total of 86 days, so the reproductive barriers caused by the phenology was 0.2711 for *E. incanum* and it was 0.0752 for *E. wilczekianum*.

**Figure 3.**
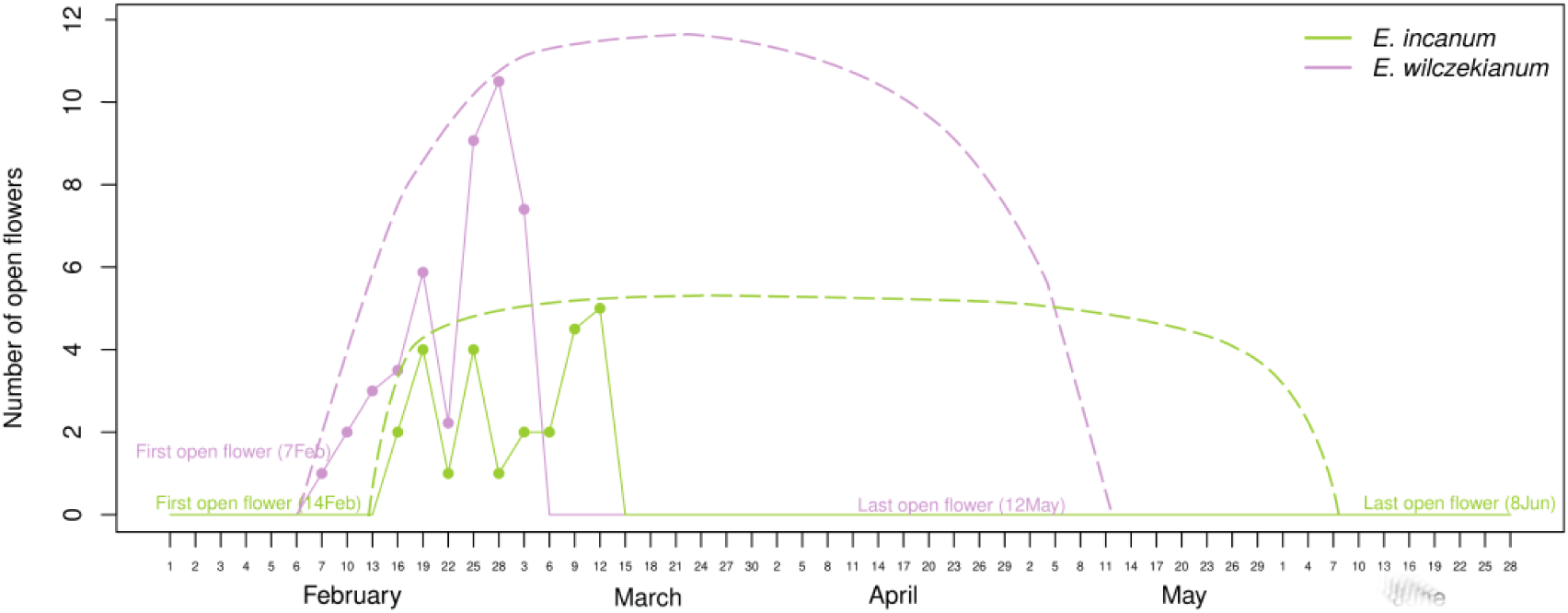
Amount of open flowers per plant shown by *E. incanum* and *E. wilczekianum* along their flowering period. Solid lines represent monitored plants during 30 days after the first open flower in *E. wilczekianum.* Dashed lines represent the inferred flowering until the last day with open flowers in each species.

### Pollinator preference isolation

A single insect was able to successively visit several flowers in the same plant of our experimental display. In fact, we noted a maximum of 20 flowers touched by a single pollinator in a plant individual. We found that *E. wilczekianum* was much more visited than *E. incanum* (254 and 4 visits, respectively, Figure 4). We also followed the pollinator flights among plants to account for intra- and inter-specific visits, being the majority of flights exclusively between *E. wilczekianum* plants. Specifically, we noted a total of 141 inter-specific flights occurred and only 2 movements were between *E. wilczekianum* and *E. incanum*. The remaining 139 flights involved flowers from *E. wilczekianum* only.

**Figure 4.**
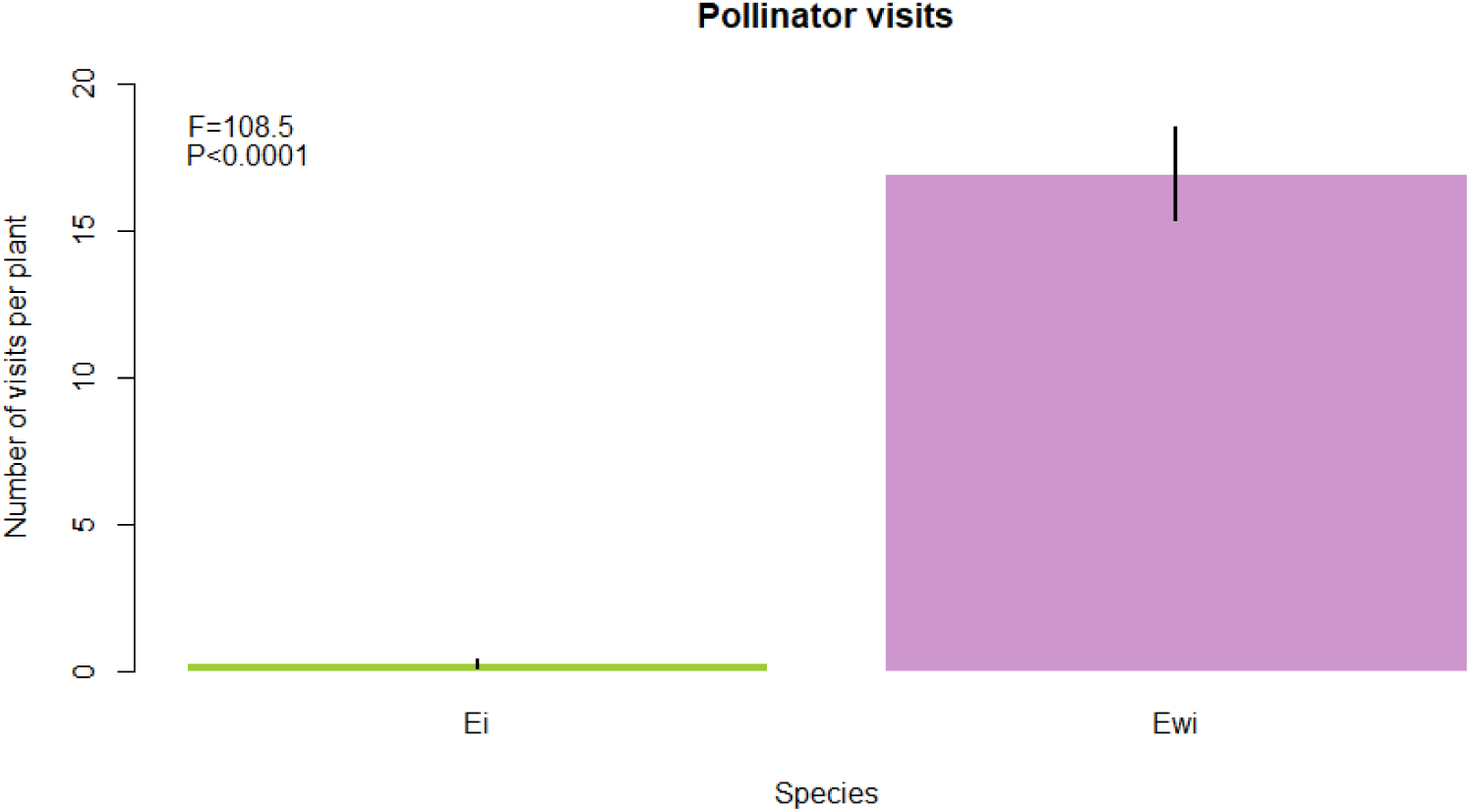
Total amount of visits received by plant individuals from *E. incanum* and *E. wilczekianum* in the two study sites.

Therefore, the reproductive barrier driven by pollinator preferences was 0.9858 for both species. In addition, we found that the frequency of visits to each species was not affected by the site where the observations were made (ANOVA; F = 0.146, *p*-value = 0.704 for the visits in *E. incanum* and F = 2.108, *p*-value = 0.150 for *E. wilczekianum* visits).

Regarding the phenotypic traits of our experimental plants, *E. wilczekianum* had larger flowers with longer corolla tubes (Fig. 5A and 5B) whereas both species showed the same amount of open flowers at the moment of the observation (Fig. 5C). Thus, insects exhibited a higher preference for visiting plants with larger corollas and longer corolla tubes (Fig. 5D and 5E) but they did not show a preference for plants showing more open flowers at that moment (Fig. 5F).

**Figure 5.**
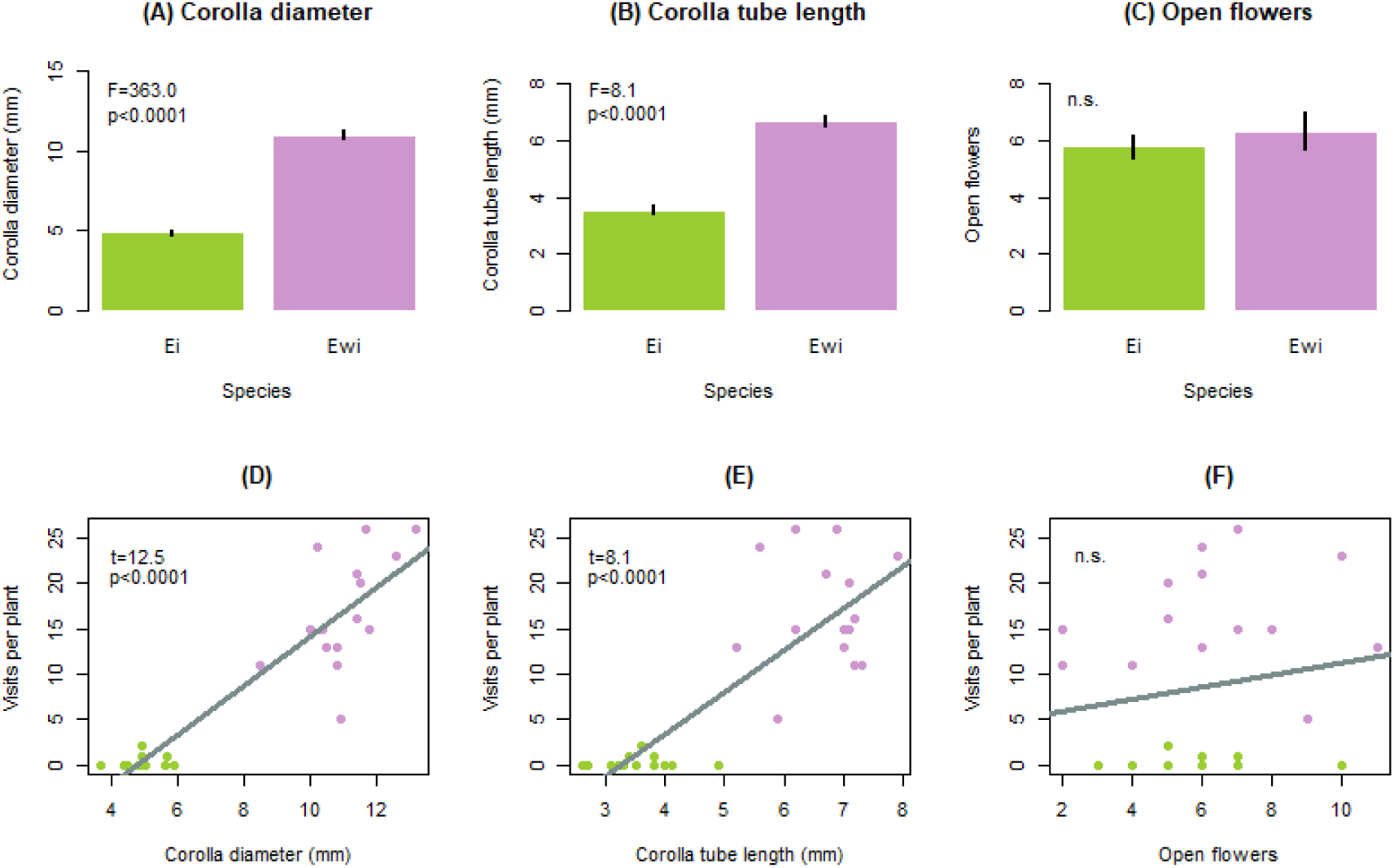
Differences in (A,B) floral size and (C) the number of open flowers prior to pollinator observations and (D,E,F) correlations between these traits and the number of visits by pollinators.

### Post-pollination reproductive barriers Hybrid seed formation

We found that the hybrid crosses when *E. incanum* was the pollen acceptor produced a high seedset, although it was lower than the seedset produced by selfing (Figure 6A; F = 110.6, *p*-value < 0.0001). In contrast, the hybrid crosses mainly failed when *E. wilczekianum* acted as the plant receptor, so we found a strong difference between the seedset from outcrossing and hybrid treatments in this species (Fig 6B; F = 637.3, *p*-value < 0.0001).

**Figure 6.**
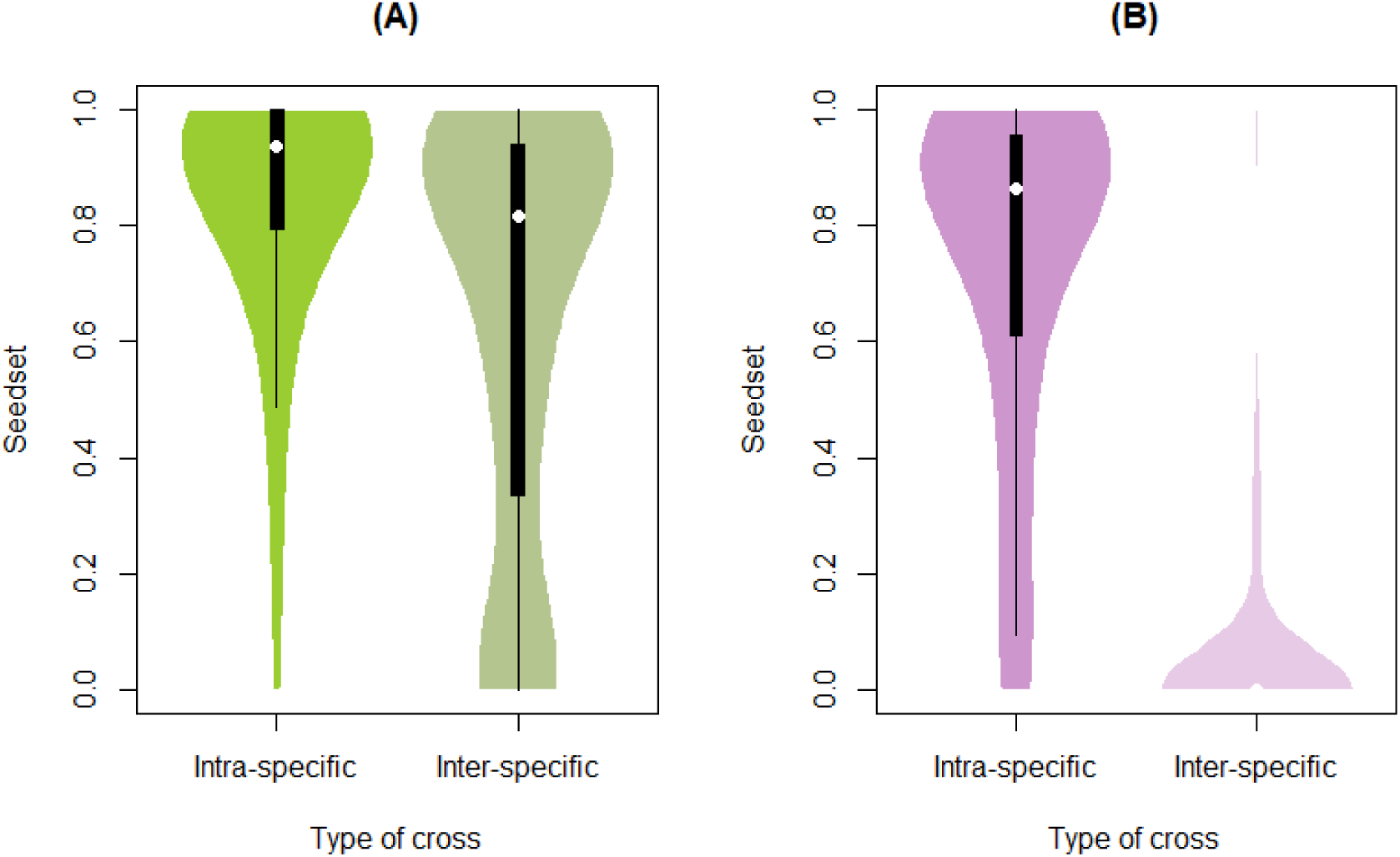
Differences in the seedset produced for (A) *E. incanum* plants through intraspecific crosses (self-pollination) and its hybrid and for (B) *E. wilczekianum* plants through interspecific crosses (cross-pollination) its hybrid.

According to these differences, we obtained a strong asymmetry in the reproductive isolation due to hybrid seed formation between both species. This reproductive barrier was 0.143 when *E. incanum* received interspecific pollen while *E. wilczekianum* frequently rejected the foreign pollen, resulting in a reproductive barrier of 0.926 in this outcrossing species.

### Viability of the first and second hybrid generation

We found a strong decrease in the germination of hybrids compared to their parental species *E. incanum* although the germination of these hybrids was not significantly different from the germination exhibited by *E. wilczekianum* which was also lower than the germination of *E. incanum* (Fig. 7A). We calculated a reproductive barrier due to differences in viability of 0.4781 between *E. incanum* and its hybrids while the reproductive barrier between these hybrids and *E. wilczekianum* would be 0.04080 due to the almost null difference in the total number of germinated plants (Fig. 7A). The germination of the seeds produced by the hybrid plants was quite similar to the germination of their parents and, thus, the same pattern of germination compared to the parental species (Fig. 7A). Hybrids from the second hybrid generation exhibit a germination significantly different from *E. incanum* but similar to both the hybrids from the first generation and *E. wilczekianum* (Fig. 7A). Therefore, reproductive barriers in this ultimate stage were similar, with the reproductive barrier due to the viability of the second hybrid generation showing a value of 0.501.

**Figure 7.**
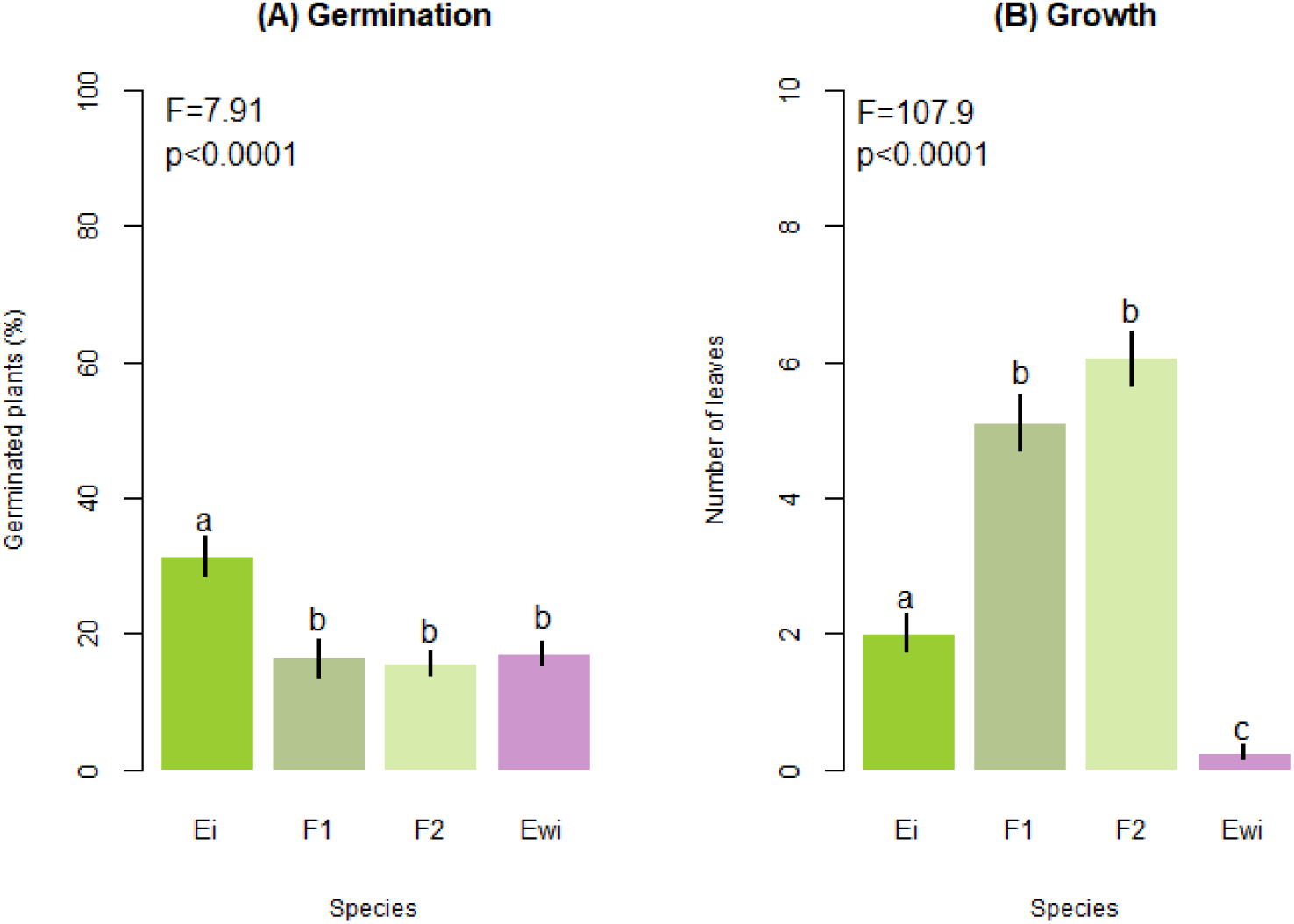
Differences in the germination proportion and the growth at first life stages among the parental species and their hybrids. Germination was considered as a viability measurement to calculate the reproductive barriers. Letters indicate significant differences. F1 and F2 refer to first and second hybrid generations, respectively.

We found that hybrids produced more leaves than both parental species at the same stage of the life cycle, indicating that hybrids grew faster (Fig. 7B). Likewise, although hybrids did not differentiate significantly from *E. wilczekianum* in germination, the leaves production was much higher. As we previously knew, *E. wilczekianum* flowers were larger (Figure 8) and produced nectar while the nectar amount in *E. incanum* was practically null (Figure 8D). We found an intermediate size in hybrid flowers compared with the parental species, although the difference between the hybrid and *E. incanum* was not significant (Fig 8). Even though the nectar amount in the first generation of hybrids was not significantly different from *E. incanum*, it tended to be increased (Figure 8D).

**Figure 8.**
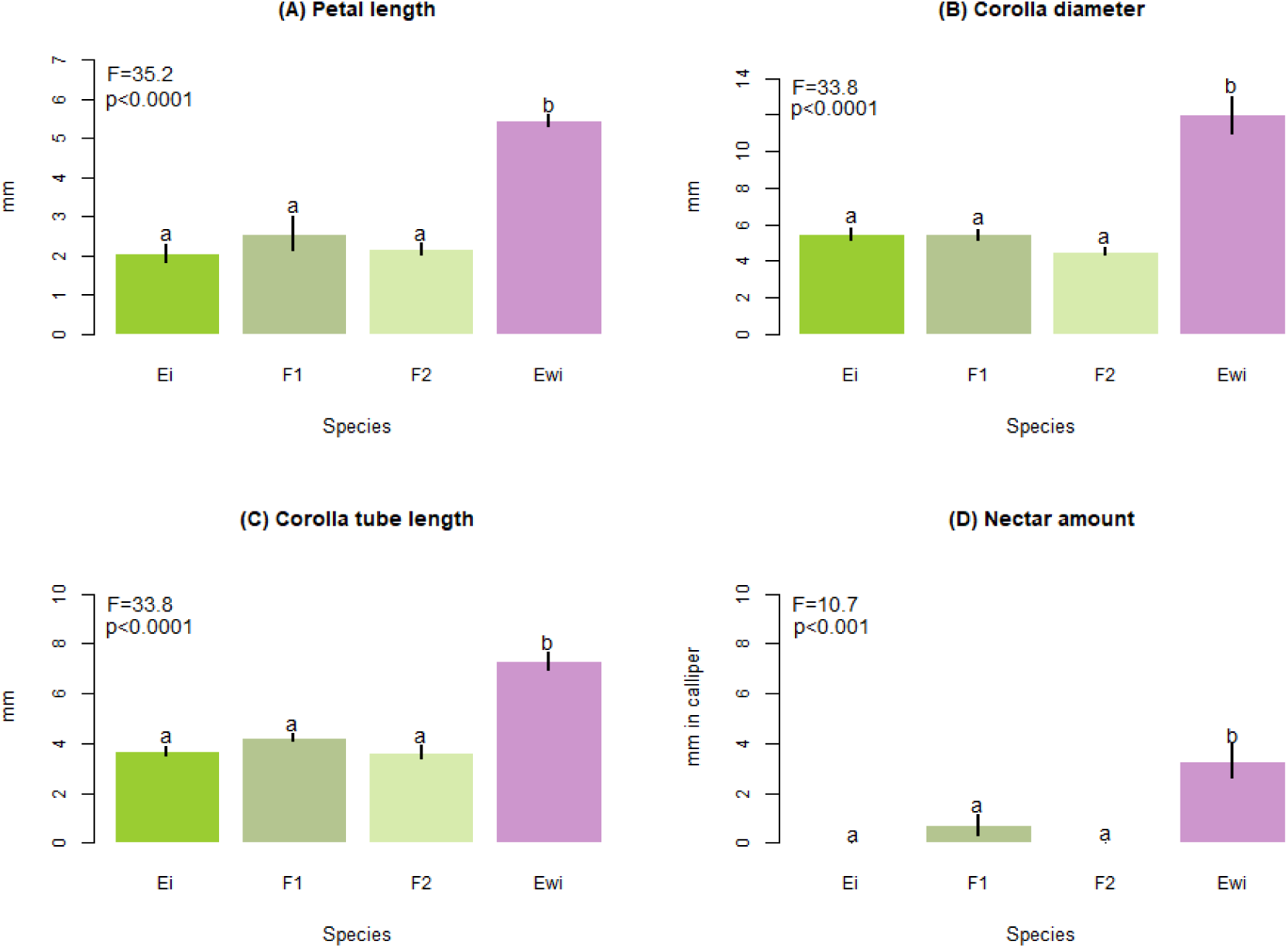
Differences in traits related to floral size as (A) the petal length, (B) the corolla diameter and (C) the corolla tube length, and (D) nectar production among *E. incanum*, *E. wilczekianum* and the first (F1) and second (F2) hybrid generation from *E. incanum*. Letters indicate significant differences.

Although the seed production of hybrid plants tended to be lower than the parental species, this difference was not significant, at least when we allowed for autonomous self-pollination (Figure 9). We compared the hybrid seedset with the seedset from *E. incanum* species to obtain a reproductive barrier due to the hybrid fertility of 0.1735.

**Figure 9.**
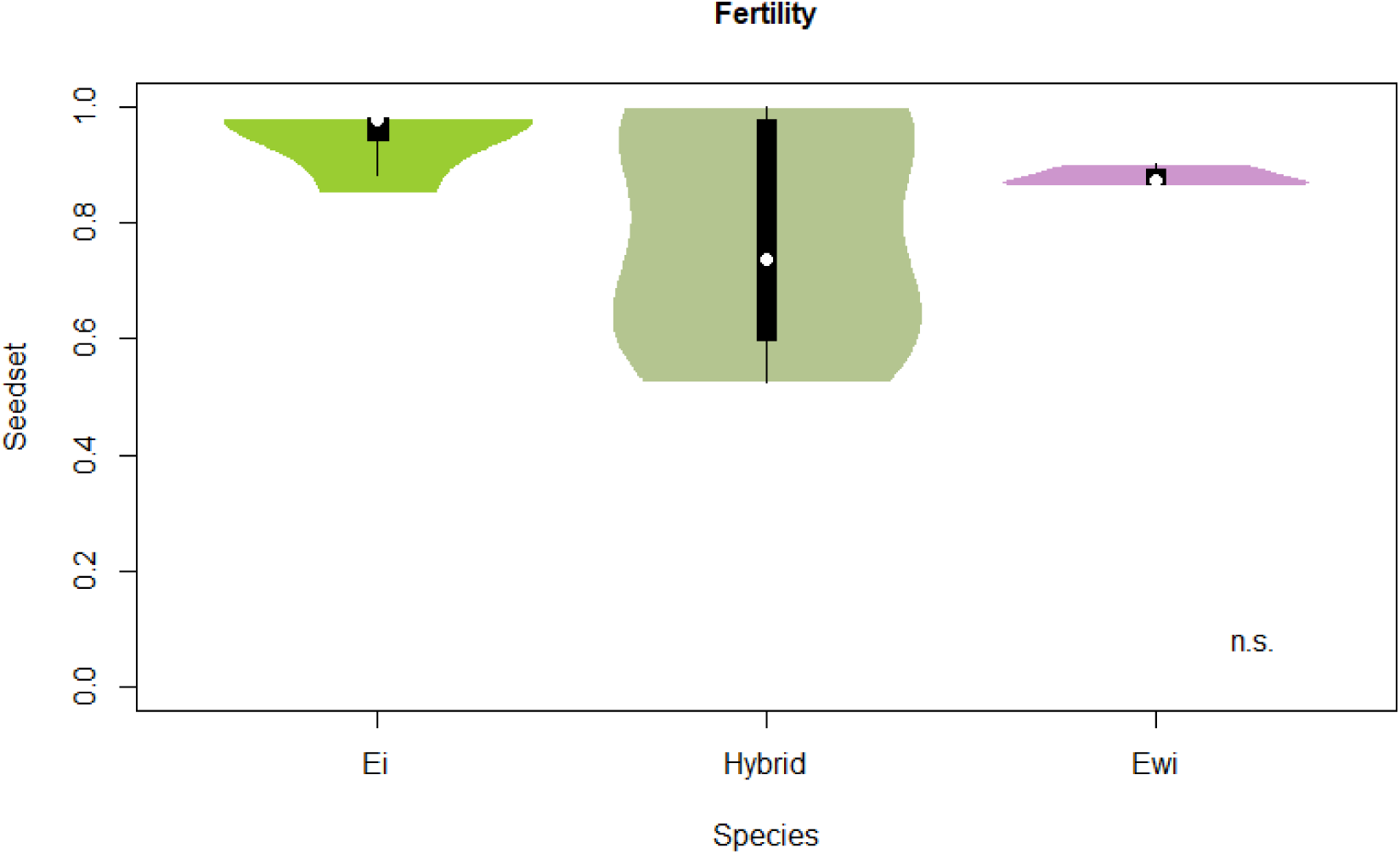
Differences in the produced seedset among *E. incanum* and *E. wilczekianum* and their hybrid from the first generation.

## DISCUSSION

The predominance of pre-pollination barriers over the comparatively weaker post-pollination barriers suggests that *E. incanum* and *E. wilczekianum* are at an early stage of ecological speciation (Nosil, 2012). This pattern has been reported in other plant systems involving recently diverged taxa (Lowry et al., 2008b; Briscoe Runquist et al., 2014), where reproductive isolation is initially driven by ecological and behavioral components before the accumulation of strong intrinsic incompatibilities. In our system, ecogeographical differentiation and pollinator preferences emerge as the main contributors to isolation, whereas phenology plays a more moderate role by reducing, but not preventing, temporal overlap in mating opportunities. Notably, the earlier flowering onset and shorter flowering duration of *E. wilczekianum* are consistent with patterns previously associated with differences in mating system strategies (Murawski and Hamrick, 1992; Stenström and Molau, 1992; Fox, 2003).

In our study system, pollinator preference appears to be the primary driver of the nearly complete reproductive isolation observed between these species. The strong bias of pollinators toward *E. wilczekianum*, characterized by larger floral displays, highlights the central role of floral traits in shaping pollinator-mediated assortative mating. In this context, floral size likely functions as a magic trait, simultaneously influencing ecological interactions and reproductive isolation (Haller et al., 2014). Importantly, these traits are closely linked to mating system evolution. The broader ecological distribution of *E. incanum* compared to the restricted niche of *E. wilczekianum* may reflect the advantages of reproductive assurance in selfing species, which facilitates colonization under conditions of limited pollinator availability (Pannell, 2015; Grossenbacher et al., 2017). Thus, differences in mating system may indirectly structure ecological divergence, reinforcing isolation through both abiotic and biotic pathways.

The contrast in mating systems between both species also extends to post-pollination processes, particularly pollen–stigma interactions. Our results show that pollen from the outcrossing *E. wilczekianum* successfully fertilizes *E. incanum*, whereas the reciprocal crosses largely fail. This asymmetry is consistent with the SI × SC rule (Murfett et al., 1996), which has been widely documented across plant taxa (Hiscock and Dickinson, 1993; Hiscock et al., 1996; Onus and Pickersgill, 2004; Baek et al., 2015). Molecular mechanisms underlying self-incompatibility are known to influence these outcomes (McClure and Franklin-Tong, 2006; Koelling and Mauricio, 2010; McClure et al., 2011), while the WISO hypothesis further predicts such asymmetries as a consequence of differential parental conflict between outcrossing and selfing lineages (Brandvain and Haig, 2005; Haig, 2000).

Beyond molecular mechanisms, ecological factors associated with mating systems may also contribute to this asymmetry. Outcrossing species such as *E. wilczekianum*, which interact with diverse pollinator assemblages, are expected to maintain more stringent pollen recognition and rejection mechanisms. In contrast, these mechanisms may be relaxed in predominantly selfing species like *E. incanum*, where the likelihood of receiving foreign pollen is reduced (Cruzan, 1990; Smith-Huerta, 1996; Kerwin and Smith-Huerta, 2000; Distefano et al., 2009). This combination of genetic and ecological factors likely explains the strong unidirectionality observed in post-pollination barriers.

The asymmetry in hybrid seed production limited our ability to fully evaluate later reproductive barriers in both crossing directions. Nevertheless, the available data indicate that hybrid performance is complex. Although hybrids exhibited reduced germination compared to *E. incanum*, their growth rate was higher than that of both parental species, suggesting partial hybrid vigour. Moreover, similar performance between first- and second-generation hybrids indicates an absence of hybrid breakdown. These results suggest that, despite initial reductions in viability, hybrid lineages may retain developmental robustness across generations.

Interestingly, hybrids also showed increased nectar production relative to *E. incanum*, representing a novel trait within this lineage. This shift may reflect the introduction of genetic variation through outcrossing and could have ecological consequences by modifying pollinator attraction. Such phenotypic changes may, in turn, influence mating system dynamics, as increased attractiveness could promote higher rates of outcrossing. Indeed, variation in selfing rates within species has been proposed as a driver of reproductive isolation and diversification (Wendt et al., 2002; Palma-Silva et al., 2015). Although the extent of hybridization in natural populations remains unknown, these results highlight the potential for hybridization to reshape reproductive strategies.

Unidirectional reproductive isolation is a common outcome in plant systems involving contrasting mating systems (Tovar-Méndez et al., 2014; Li and Chetelat, 2015). Experimental studies in other taxa, such as *Mimulus*, have shown that interactions between floral traits and mating systems can strongly influence the strength and direction of reproductive barriers (Martin and Willis, 2007). Our findings are consistent with this framework, as the traits contributing most to reproductive isolation are directly linked to the reproductive strategies of each species.

The role of mating system as a reproductive barrier has been widely debated (Coyne and Orr, 1998; Fishman and Stratton, 2004; Lowe and Abbott, 2004; Martin and Willis, 2007). While mating system per se may not always be considered an independent barrier, it can shape multiple mechanisms that collectively reduce gene flow. In this study, we argue that the selfing strategy of *E. incanum* acts as an effective barrier by limiting pollinator-mediated gene flow through the expression of the selfing syndrome. Traits associated with selfing, such as reduced floral attractiveness, directly constrain interactions with pollinators and thereby reinforce ecological isolation. In this sense, mating system should be viewed not as a single barrier but as a central axis structuring multiple components of reproductive isolation.

Taken together, our results contribute to a broader understanding of how mating system divergence can shape the architecture of reproductive isolation in plants. Rather than acting as a single barrier, mating system differences appear to reorganize multiple components of isolation simultaneously, from pollinator-mediated processes to post-pollination compatibility. This integrated perspective is particularly relevant in the context of evolutionary transitions, where shifts in mating strategy may trigger coordinated changes across ecological and reproductive dimensions. More generally, our findings highlight the importance of evaluating reproductive isolation as a composite and dynamic process, especially in systems where gene flow is still possible. By linking ecological divergence, mating system evolution, and the accumulation of reproductive barriers, this study provides a framework for understanding how complex interactions among traits can drive speciation in flowering plants.

## ACKNOWLEDGEMENTS

We thank all the members at the research network BioChange (http://biochangenet.org) for their lab assistance and paper discussion.

## AUTHOR CONTRIBUTIONS

MA and AJMP conceived the initial idea of this study, which was further developed by AGM, CF, EO. AGM, CF, EO and MA performed the manual controlled crosses and evaluated the plant fitness. All the authors contributed to evaluate the reproductive barriers and perform the statistical analyses, leaded by AJMP. AGM wrote the first draft of the manuscript. All authors contributed to the interpretation of the results, as well as to the revision and improvement of the final version of the manuscript.

## DATA AVAILABILITY

The data that support the findings of this study are available in the supplementary material and the corresponding author upon reasonable request.

## SUPPLEMENTARY MATERIAL

**Figure S1.**
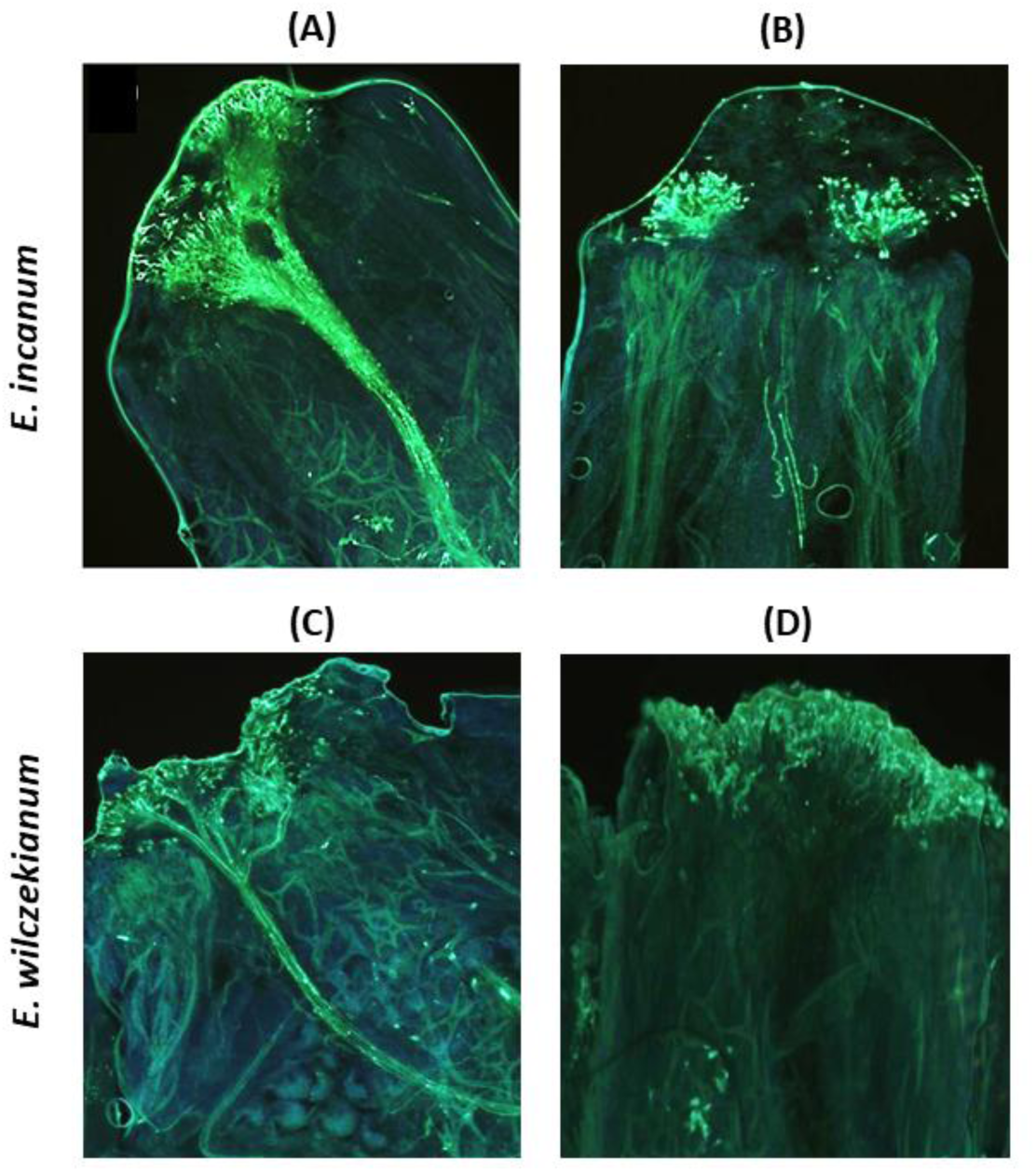
Visualization of the interaction between pollen and stigma observing the growth of pollen tubes by UV luminescence for: (A) intraspecific and (B) interspecific cross in *E. incanum*; (C) intraspecific and (D) interspecific cross in *E. wilczekianum*.

